# Field evaluation of a volatile pyrethroid spatial repellent and etofenprox-treated clothing for outdoor protection against forest malaria vectors in Cambodia

**DOI:** 10.1101/2024.01.30.577940

**Authors:** Élodie A Vajda, Amanda Ross, Dyna Doum, Emma Fairbanks, Nakul Chitnis, Jeffrey Hii, Sarah J Moore, Jason Richardson, Michael Macdonald, Siv Sovannaroth, Pen Kimheng, David J McIver, Allison Tatarsky, Neil F Lobo

**Affiliations:** University of California, San Francisco, 550 16th Street, San Francisco, CA, 94158, USA; Swiss Tropical and Public Health Institute, Socinstrasse 57, PO Box, CH-4002, Basel, Switzerland; University of Basel, Petersplatz 1, CH-2003, Basel, Switzerland; Health Forefront Organization, Phnom Penh, Cambodia; Vector Control Product Testing Unit, Environmental Health and Ecological Science Department, Ifakara Health Institute, P. O. Box 74, Bagamoyo, Tanzania; Innovative Vector Control Consortium, Liverpool School of Tropical Medicine, Pembroke Place, Liverpool, Merseyside, L3 5QA; National Center for Parasitology, Entomology and Malaria Control, 477, Phnom Penh, Cambodia; Department of Health of Mondulkiri, C5XX+CP4, 76, Krong Saen Monourom, Cambodia; University of Notre Dame, Notre Dame, IN, 46556, USA

## Abstract

Cambodia’s goal to eliminate malaria by 2025 is challenged by persisting transmission in the country’s forest and forest fringe areas. People living in, or traveling to the forest, are exposed to malaria vector bites during the day due to *Anopheles* daytime biting; and during the night, due to low bed net use and open sleeping structures. Volatile pyrethroid spatial repellents (VPSRs), and insecticide treated clothing (ITC) may help address these gaps in protection. In this field study the authors evaluated the outdoor application of one passive, transfluthrin-based VPSR, four etofenprox-ITCs paired with a picaridin topical repellent, and a combination of VPSR and ITC against wild *Anopheles* landing in Cambodia. Mathematical modeling was also used to predict the reduction of vectorial capacity of these interventions.

A 7×7 Latin-square (6 interventions and one control) was conducted over 49 collection nights in seven temporary, open structures in a forest in Mondulkiri Province, Cambodia. Pairs of participants conducted human landing catches (HLCs) from 18h00 to 06h00, with each collector conducting collections for six hours. A randomly selected subset of collected *Anopheles* were identified to species using molecular methods. The rate ratio of each intervention compared to the control on *Anopheles* landings was estimated using a mixed-effect negative binomial regression with intervention, structure, and collector-pair as fixed-effects, and with collection date and structure-night as random effects. The modeling assessment aims to predict the relative reduction in vectoral capacity. Initial calculations involved establishing a “baseline scenario” without intervention, utilizing biometric parameters for *Anopheles dirus*. Various scenarios accounting for intervention coverage and adherence were then considered. The study aims to update parameters using field study estimates for wild *Anopheles*, incorporating multiple semi-field estimates for interventions and accounting for the variability and uncertainty in parameter values.

Of the total 8,294 *Anopheles* specimens collected, 15% (n=1,242) of specimens were confirmed to species or species group via PCR. Fifteen species were confirmed; *Anopheles dirus* Form A was predominant (n=429), followed by *Anopheles maculatus* (n=189), and *Anopheles minimus* (n=60). All six interventions reduced *Anopheles* landing substantially; protective efficacies ranged between 61% (95% confidence interval (CI): 48 – 71%) (etofenprox-ITC, washed) and 95% (95% CI: 93 – 96%) (combined VPSR and unwashed etofenprox-ITC). Finally, the modelling assessment demonstrates significant reductions in vectoral capacity, with the highest impact observed for the combined ITC and VPSR as well as the VPSR used alone, although effectiveness decreases with intervention aging, and variability exists in the magnitude of predicted reductions due to differences in experimental conditions.

**T**hese transfluthrin-based VPSR and etofenprox ITC interventions have the potential to reduce outdoor and daytime *Anopheles* biting by providing substantial protection against *Anopheles* landing. One or more of these tools may play a valuable role in the push for elimination in Cambodia and the Greater Mekong Subregion if programs can achieve effective coverage.

## Introduction

The Greater Mekong Subregion (GMS) aims to achieve elimination of *Plasmodium falciparum* malaria by 2025, and of all human malaria by 2030 [1, 2]. From 2000 to 2020, the GMS recorded a 56% decrease in malaria, and an 89% reduction in *P. falciparum* cases. As of 2022, Cambodia accounts for 2% of all malaria cases and for 1% of *P. falciparum* cases in the GMS. Operating on an accelerated timeline compared to the remaining GMS countries, Cambodia aims to eliminate *P. falciparum* malaria by 2024, and all human malaria by 2025 [1].

From 2010 to 2022, confirmed malaria cases in Cambodia declined by 91%. As malaria decreases throughout the country, it has become increasingly confined to malaria transmission foci [2]. Roughly 0.5 million people in Cambodia live in forest and forest fringe areas characterized by high malaria transmission [3,4]. These forest transmission foci are predominantly inhabited by ethnic minorities, local populations, and rural mobile and migrant populations working in rubber plantations, mining, and agriculture [5–7]. Under the Malaria Elimination Action Framework (MEAF) (2021-2025), the Cambodia’s National Center for Parasitology, Entomology, and Malaria Control (CNM) deploys forest packs, often containing insecticide-treated nets (ITNs), insecticide treated hammock nets (ITHNs), and topical repellents to mobile and migrant populations in areas at highest malaria risk [8,2]. Since November 2020, Cambodia has also adopted foci-based innovative strategies as part of its “last mile” strategy towards *P. falciparum* malaria elimination, including targeted drug administration (TDA) and intermittent preventive treatment for forest-goers (ITPf) [9,10].

Appropriate bite prevention interventions to reduce forest-going and -dwelling populations’ exposure to forest-based *Anopheles* are needed based on the spaces and times where, and when these individuals are exposed to *Anopheles* [4]. A recent study conducted in northern Cambodia found that while *Anopheles* densities strongly declined in villages during the dry season, densities remained relatively similar from the dry to the wet season in the forest [7]. Therefore, forest sites may serve as suitable refuges for *Anopheles* during the dry season, and consequently, also as a malaria parasite reservoir, since human activity in the forest is particularly extensive during the dry season [7]. In addition, forest-goers are exposed to vector bites during the day due to *Anopheles* exhibiting outdoor, daytime and early evening biting [11]; and during the night, due to low bed net use and open sleeping structures [7,12]. These characteristics limit the effectiveness of strategies that focus on traditional village- and homestead-centric vector control interventions (indoor residual spraying (IRS), insecticide treated nets (ITNs)) [6,13].

Additional interventions that target mosquitoes and people outdoors must be available for use alongside IRS and ITNs. Two promising tools have been developed: volatile pyrethroid spatial repellents (VPSRs), and insecticide treated clothing (ITCs). VPSRs function by preventing human-vector contact primarily through non-contact irritancy (also referred to as non-contact excito-repellency or spatial repellency), landing inhibition, feeding inhibition, sublethal incapacitation, and pre-/post-prandial (before/after blood feeding) mortality [14]. ITCs primarily protect humans from mosquito bites through contact irritancy (also referred to as contact excito-repellency), some short-range non-contact excito-repellency, feeding inhibition, and mortality [15–17]. VPSRs have been extensively evaluated in Africa [18– 24] and increasingly in Southeast Asia [25–29]. ITCs, treated with pyrethroids, have been widely evaluated and show promise for their use against *Anopheles* biting amongst mobile populations and military/ranger personnel [30–33]. However, commonly used permethrin formulations have shown poor wash retention and low bite prevention levels on many lighter weight fabrics prompting the development of etofenprox clothing treatment products with superior wash retention and safety profiles [34,35]. The evidence that permethrin treated clothing provides protection against malaria is unclear [16,36–38] and the WHO currently does not recommend its use for the prevention and reduction of malaria at the community level where malaria transmission is ongoing [39].

As part of a phased approach to vector control product evaluation [40], a series of preceding semi-field system (SFS) experiments were conducted in Thailand to measure the primary effects (landing prevention as a proxy for bite prevention) and the secondary effects (sublethal incapacitation and mortality) of new transfluthrin- and metofluthrin-based VPSRs, and etofenprox-treated clothing, against pyrethroid-susceptible *Anopheles minimus* (an important vector of malaria in the GMS [41]). In addition to providing insights and estimates of intervention impact on landing inhibition, sublethal incapacitation, and mortality, the data were used to parameterize a mathematical model to predict the potential of the tools to provide community protection [42]. Vectoral capacity is a measure of the ability of the vector population to transmit a disease, defined as the total number of potentially infectious bites that would eventually arise from all the mosquitoes biting a single infectious human on a single day [43]. Interventions that reduce vectoral capacity have the potential to confer community-wide protection leading to reduced malaria burden [44].

Building on the SFS studies, an entomological field study was undertaken to evaluate these VPSR and ITC interventions on wild *Anopheles* landing using human landing catches (HLCs) in a small-scale, controlled field setting in Cambodia. The study’s secondary objectives were to 1) validate the results from the preceding SFS evaluation, 2) measure the hourly human landing rates (HLR) in the control structures, towards describing the local *Anopheles’* evening and nighttime biting trends, 3) confirm the *Anopheles* species identification from a subset of the HLC-collected specimens, and to 4) predict the reduction of vectorial capacity of these interventions using mathematical modelling. For objective 4, the potential of these tools to provide protection beyond personal protection is investigated by combining data from both the field study in Cambodia and the preceding semi-field studies in Thailand.

## Methods

### Study site

Field collections took place in the village of Andong Krolong (12.320725, 107.029779), Mondulkiri Province, Cambodia. The study took place from 23 September to 24 November 2022, thus taking place during the rainy season and the early days of the dry season. Temporary, open sleeping structures representative of open sleeping structures used by locals were constructed out of wooden poles and tarpaulin roofs (2×2×2 m) (Fig 1). While closely surrounded by forest cover, the open structures were in an area that had been recently cleared, representative of the living conditions of forest workers (Fig. 1).

**Fig. 1.**
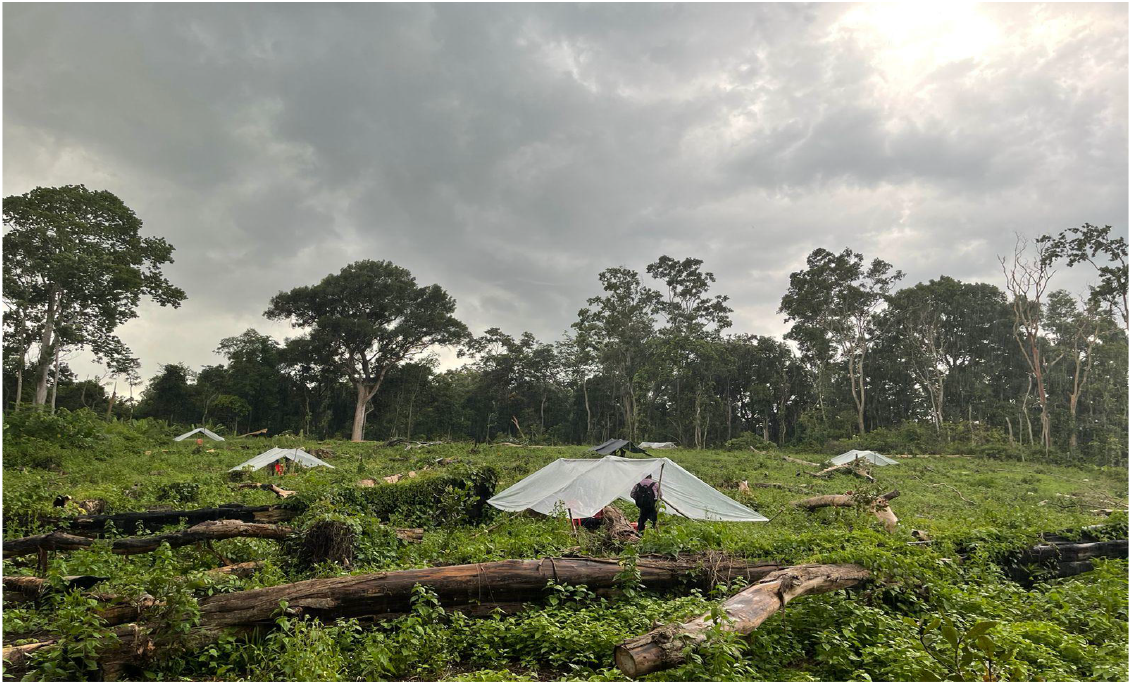
Temporary, open structures set-up in the field site in Andong Krolong, Mondulkiri Province, Cambodia.

### Mosquito bite prevention interventions

Six bite prevention interventions were evaluated (Table 1). Interventions included a transfluthrin-based VPSR (BiteBarrier (‘BB’), formerly known as PIRK), civilian and ranger clothing that were either newly treated with etofenprox or treated with etofenprox and then washed 20 times, as well as one arm with both the VPSR and etofenprox-treated civilian clothing. The treated civilian and ranger clothing was always paired with a 20% picaridin topical repellent (*SC Johnson*). The picaridin repellent was selected for this study as it has previously been demonstrated to be safe and effective against Southeast Asian vectors of malaria [11].

**Table 1.**
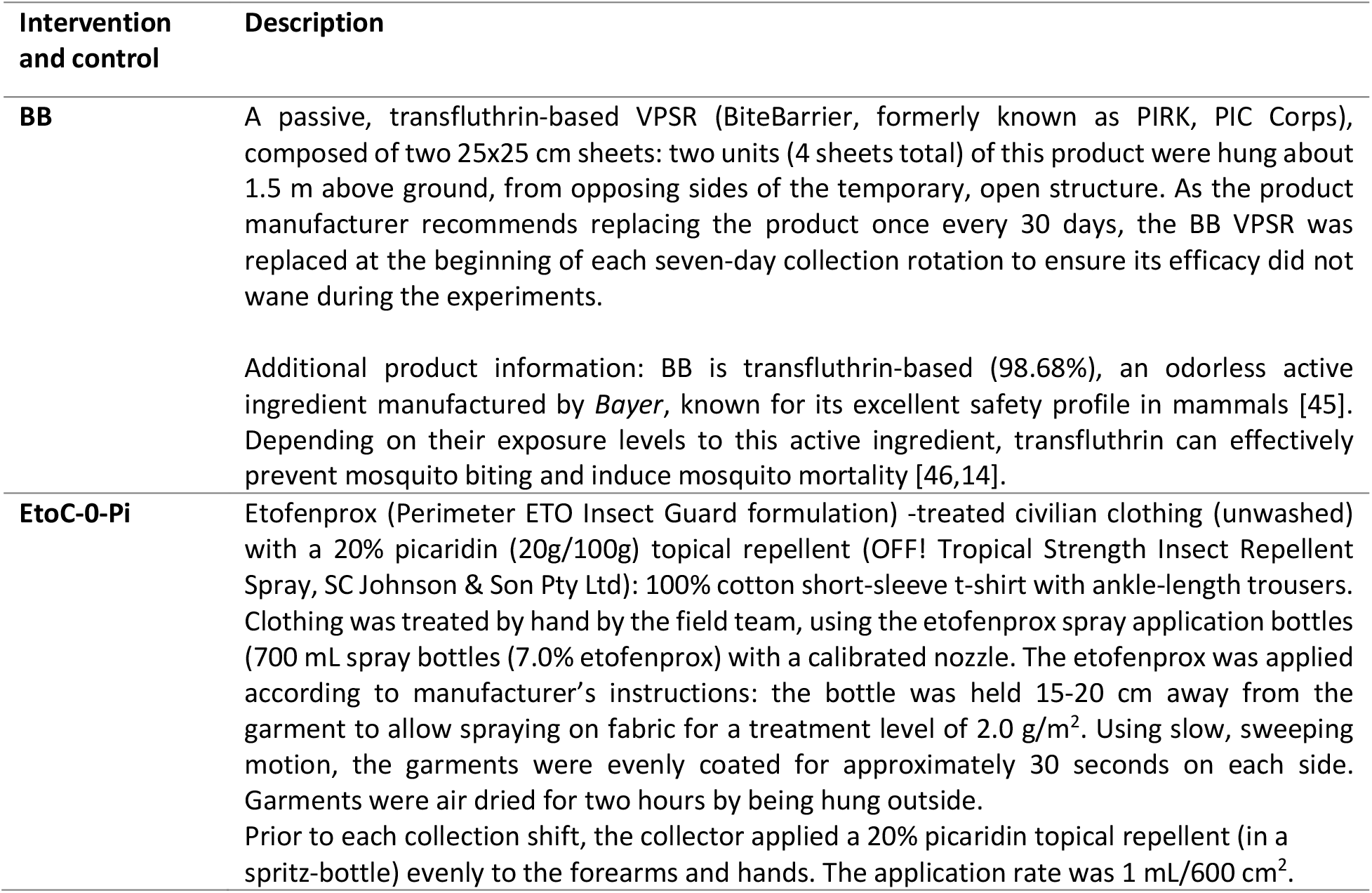
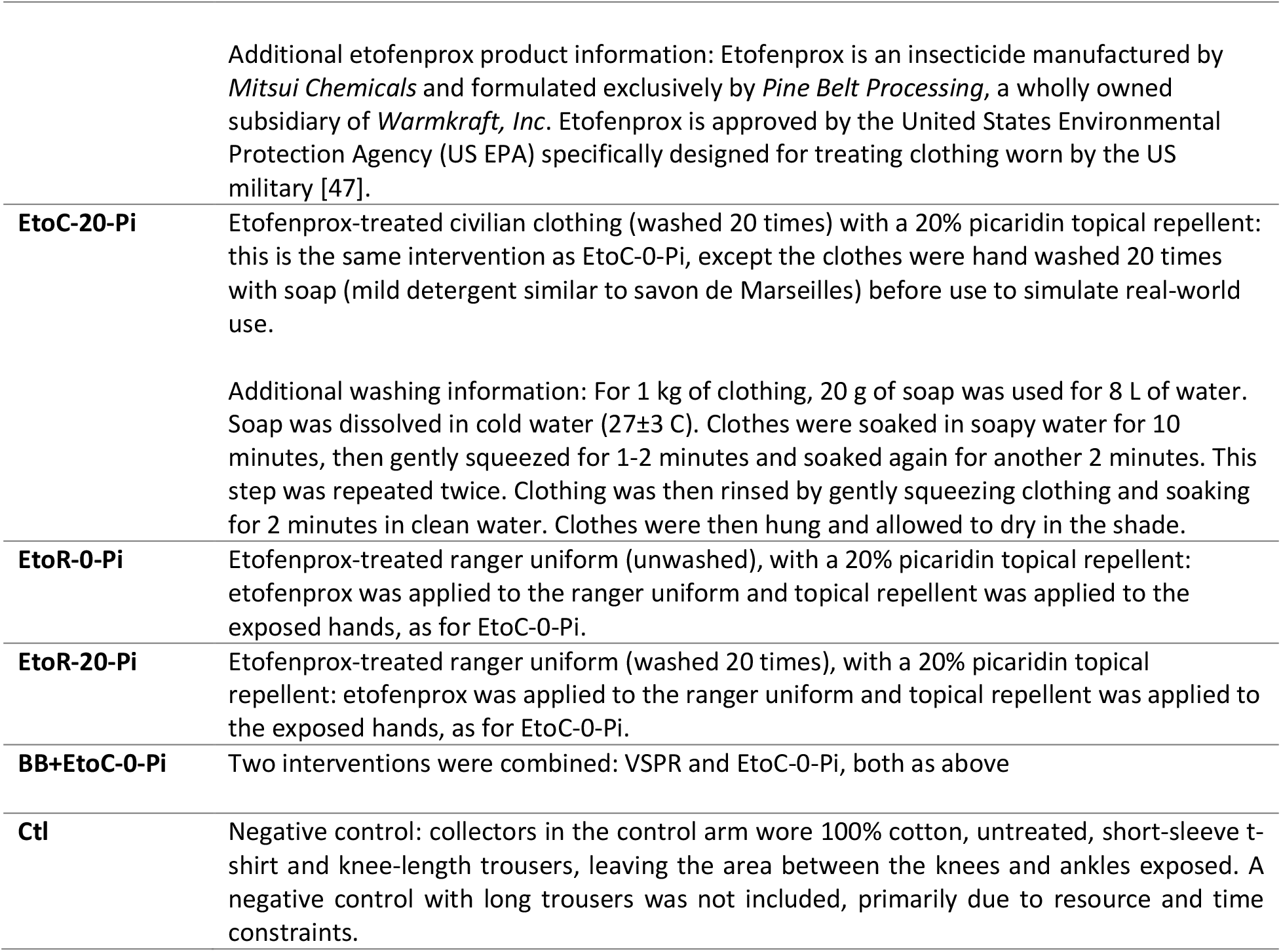
Descriptions of the mosquito bite prevention interventions and the control.

### Experimental design and mosquito collections

Seven temporary, open structures located at least 20 m apart were set up. Human landing catches (HLCs) were carried out in the structures for 12 hours, from 18h00 to 06h00, divided into two collection shifts, 18h00 to 00h00 (shift 1) and 00h00 – 06h00 (shift 2). A fully balanced 7 × 7 Latin-square design was used. Each of the seven study arms (six interventions and one control) was assigned to one structure for seven collection nights, and each pair of collectors rotated through each location on a nightly basis. After each block of 7 collection nights, interventions were advanced to the next position, and collectors continued to rotate through structures each night. After 49 nights of collection each collector had tested each intervention in each location seven times. There were 20 unique HLC collector pairs. Some collector pairs worked fewer HLC nights than others as some individual collectors left the study before its completion. Due to cultural perceptions about being in the forest at night, collector pairs remained together in the structure throughout the entire collection night: while one collector worked, the other collector slept in an untreated hammock net.

HLCs were used to collect mosquitoes landing on the area from knee to ankle of the collector for each collection rotation. All HLC collectors provided written informed consent and were provided free-of-cost malaria diagnosis and treatment should malaria symptoms (e.g., fever) occur during the study and/or during the two weeks subsequent to the field trial period. For etofenprox-treated clothing interventions, long trousers were not rolled up to the knee in order to estimate the landing protection afforded by the combination of etofenprox with long trousers. The negative control had the area between the knee and the ankle exposed. The total number of mosquitoes caught hourly was recorded. Mosquitoes captured were stored in individually labelled Eppendorf tubes (by treatment and hour of collection), transported in coolers to the base camp every morning, killed by freezing, counted, morphologically sorted, and stored individually with desiccant in Eppendorf tubes for subsequent processing.

### Environmental data collection

Temperature (Celsius), relative humidity (%RH), windspeed (m/s) and rainfall occurrence were recorded on an hourly basis during HLC collections, from 18h00 to 06h00. Temperature, %RH, and wind speed data were recorded at the end of each HLC collection hour using a data logger device (HOBO®). Rainfall occurrence was recorded as a binary variable (yes/no) to indicate occurrence or absence of rainfall during each HLC collection hour.

### Mosquito morphological and molecular species identification

All *Anopheles* mosquitoes were sorted to species or species group using morphological identification keys [48] in the field. Individual specimens were then packaged in individual, tightly closed, Eppendorf tubes with silica gel and were sent to University of Notre Dame (UND), Indiana, USA, for molecular species confirmation. Approximately 15% of samples were randomly selected across all HLC collections, and sequenced at the ribosomal DNA internal transcribed spacer region 2 (ITS2) and/or cytochrome oxidase subunit 1 (CO1) loci towards species determination [12]. Conservative molecular species identification was based on matches to GenBank (National Center for Biotechnology Information [NCBI]) and BOLD [49] (databases with lower quality matches and an absence of voucher specimens resulted in identifications to higher taxonomic levels).

### Data analysis

The change in landing associated with each intervention was estimated as rate ratios (RR), the ratio of the number of *Anopheles* landing in the intervention compared to that in the control. They were estimated using a mixed-effect negative binomial regression with structure ID, collector pair, and intervention ID as fixed-effects, and with collection date and batch (the location-night) as random effects. The effect of structure location and collector pair on *Anopheles* catches per collection night was also examined. All RR estimates are presented with 95% confidence intervals (CI). The percentage protective efficacies of each intervention against *Anopheles* landing were estimated as (1−*RR*)×100.

The relationship between weather variables and *Anopheles* densities caught was investigated. Mean nightly temperatures, mean nightly %RH, mean nightly wind speed, and number of rain occurrences per night were plotted against the total number of *Anopheles* captured per night. All statistical analyses were performed in R (Murray Hill, New Jersey), Version 2023.03.0+386 [50], using the tidyverse packages ‘tidyr’ [51], ‘dplyr’ [52], ‘lme4’ [53], and ‘ggplot2’[54].

The hourly human landing rate (HLR) for the control arm across the 49 collection nights was estimated as a proxy for the hourly human biting rate (HBR) [55,56].

### Modeling of vectoral capacity

The relative reduction in vectoral capacity was predicted using the method described in Denz et al (2021) [44] and in Fairbanks et al (preprint, 2023) [42]. Firstly, the vectoral capacity is calculated for a “baseline scenario” (without the intervention). Parameters used in the calculation for this are given in Suppl Table

1. Biometric parameters for *Anopheles dirus* were used since this species accounted for most of the local mosquito population. Next, were considered other scenarios where a proportion of the population is protected by an intervention. This proportion is dependent on both the coverage (whether someone has the intervention) and adherence (whether someone uses the intervention).

Fairbanks et al (preprint, 2023) [42] used data from semi-field studies combined with a model to quantify the modes of action of selected vector control tools. The influence of the interventions on the mosquito feeding cycle were described relative to an unprotected human host using four characteristics; the relative reduction in the rate of landing, the relative increase in the rate of preprandial killing and disarming, the proportion of this increase in rate which is due to disarming, and the change in the probability of postprandial mortality. We will use estimates from the field study to update some of these parameters to provide estimates for the wild *Anopheles* at this field site. For some interventions there are multiple semi-field estimates (from different sites and years), in this case we consider estimations from each semi-field scenario separately, and therefore have multiple estimates for the reduction in vectorial capacity for these interventions.

The relative reduction in the rate of biting parameters estimated from the field study were used. Then, to consider the increase in the rate of preprandial killing and disarming, we first calculate the relative change in the rate of biting in the field studies compared to semi-field studies:

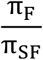

where π_*F*_ and π_*SF*_ are the median estimated protective efficacies for the change in the rate of biting in in the field and semi-field studies, respectively. This ratio helps us understand how the reduction in biting in the field studies compares to the semi-field studies. The reduction in the rate of disarming and preprandial killing is then calculated as:

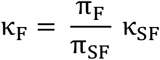

where *κ*_*SF*_ is the estimated rate from the semi-field data. This scaling method ensures that the proportion of the reduction in biting attributed to disarming or preprandial killing matches that observed in the semi-field studies, while the magnitude of these effects are assumed to scale according to observation under field conditions (with wild *Anopheles*). The probability a mosquito is killed postprandially and the probability a mosquito is disarmed, given it is disarmed or killed preprandially, are assumed to be the same as in the semi-field conditions.

To incorporate variability and uncertainty, parameter values are sampled from the protective efficacy distribution (reduction in biting) or the from the semi-field study (all other parameters).

## Results

### Anopheles species identification and human landing rates

A total of 8,294 *Anopheles* specimens were collected with HLCs. Of this total, 96% (n=7,951) of specimens were morphologically identified to species or species group. Out of the total morphologically identified, 96% (n=7,621) were identified as *An. dirus* sl, leaving the remaining 5% of specimens identified to eight different species or species group: *Anopheles maculatus* sl (n=234), *An. minimus* (n=63), *Anopheles philippinensis* (n=10), *Anopheles kochi* (n=8), *Anopheles aconitus* (n=7), *Anopheles barbirostris* sl (n=5), *Anopheles baimaii* (n=2), and *Anopheles asiaticus* (n=1). Molecular speciation of 15% (n=1,242) specimens were evaluated, and confirmed that the morphological identification was extremely accurate (Table 2).

**Table 2.**
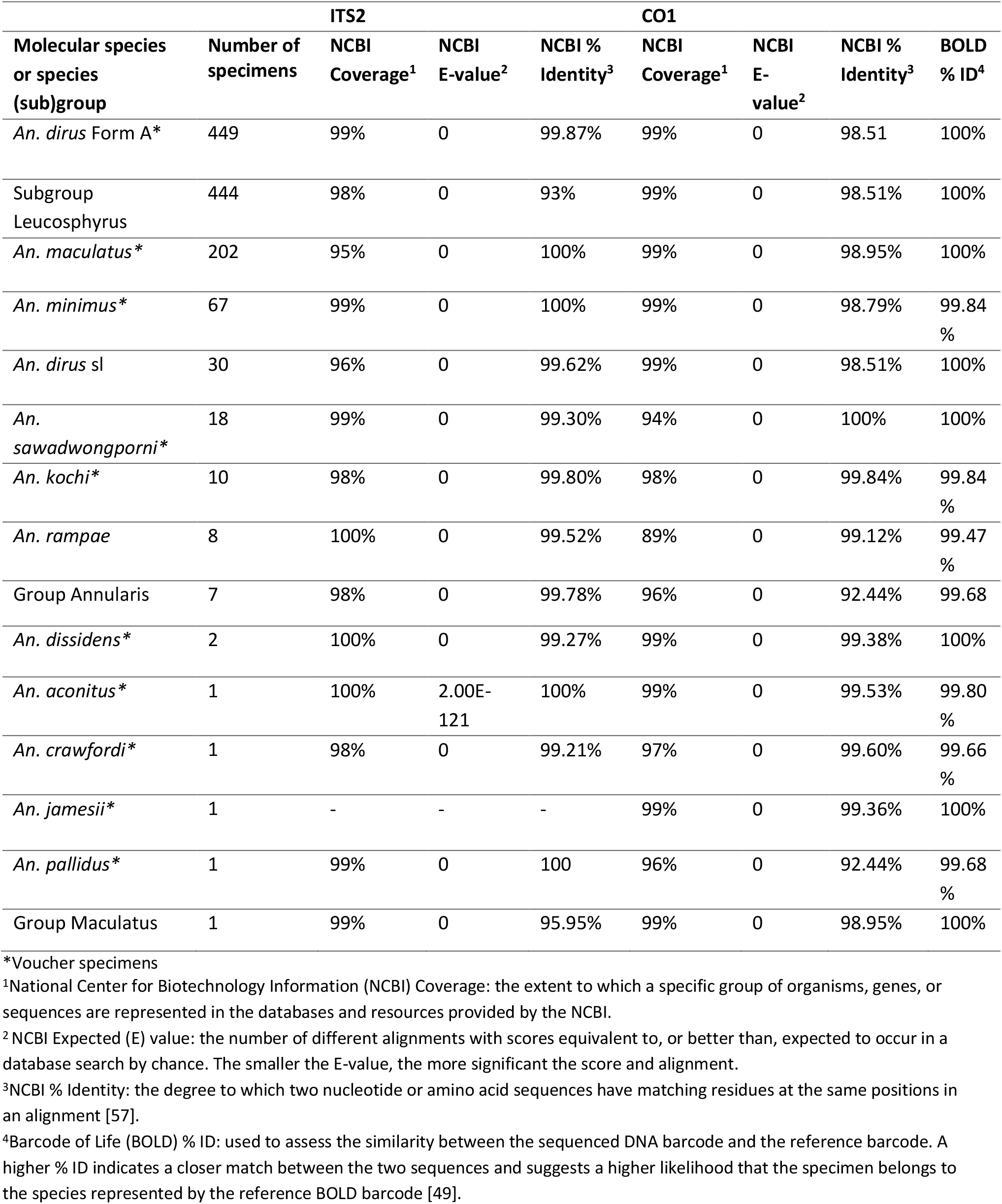
*Anopheles* species and species group confirmed via molecular species identification.

The estimated mean HLR per hour (control arms) ranged from 8.16 (95%CI: 0.21 – 16.10) *Anopheles* landings per person per hour (lph) (00h00 – 01h00) to 2.61 (CI: 0.00 – 5.22) lph (05h00 -06h00). Higher landing rates were recorded from 18h00 to 00h00. After 02h00, a decline of the mean hourly HLR was observed until the last HLC collection hour (05h00 – 06h00) (Fig 2).

**Fig. 2.**
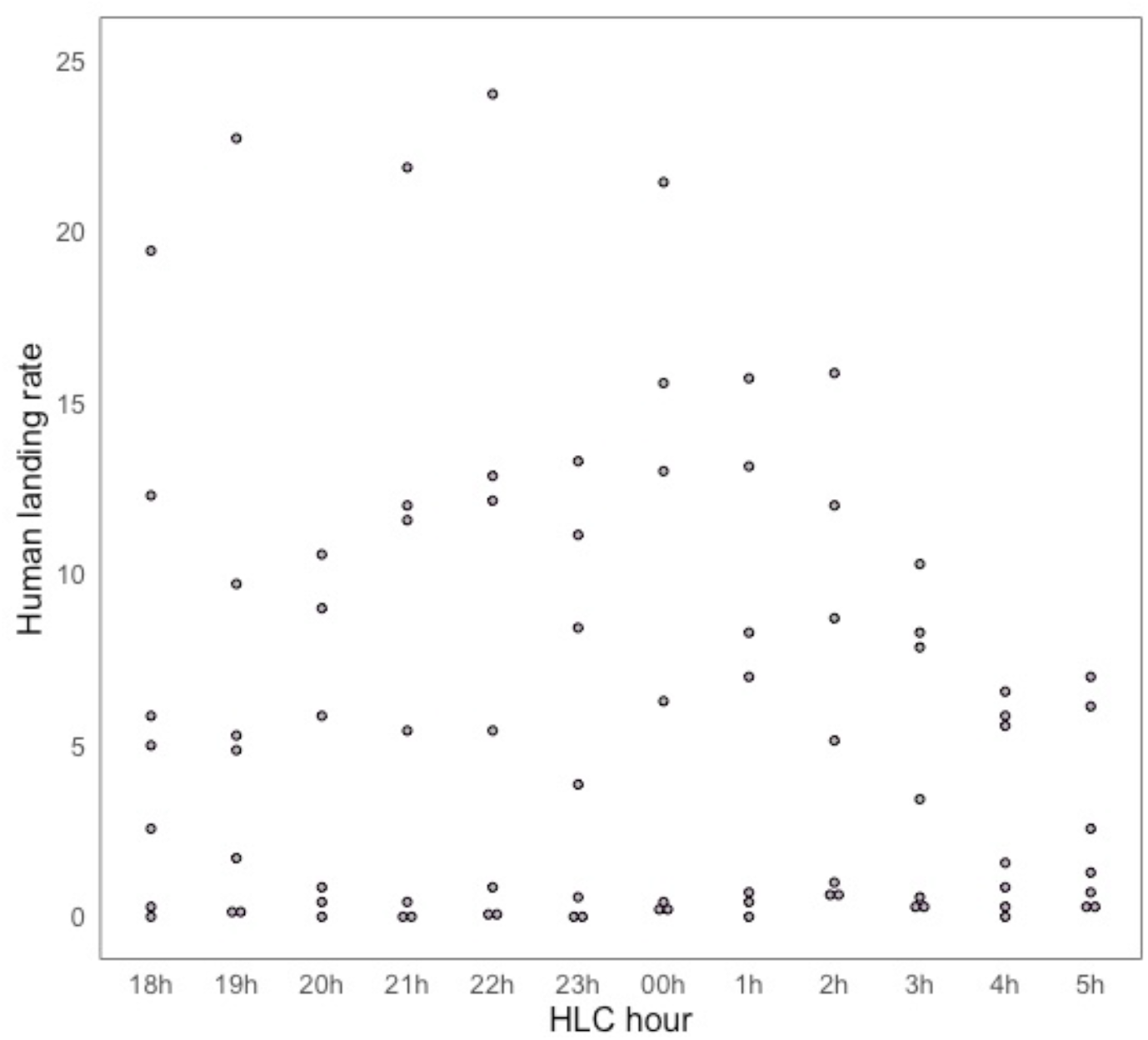
Hourly *Anopheles* human landing rate (HLR) (*Anopheles* landings per person, per hour), over the course of 49 collection nights (from control arms).

### *Intervention protective efficacy against* Anopheles *landing*

All six mosquito bite prevention interventions showed substantial reductions in mosquito landings relative to the control (Ctl) (Fig. 3). The estimated protective efficacies of the interventions against *Anopheles* landing ranged from 61% (95%CI: 48 - 71%) to 95% (CI 93 - 96%) (Fig. 4, Table 3). BB alone and the combined interventions (BB+EtoC-0-Pi) provided the greatest protection against *Anopheles* landing, with protective efficacies of 94% (CI 91 - 95%) and 95% (CI 93 - 96%), respectively. The etofenprox-treated ranger uniform and civilian clothing, unwashed, provided similar protection against landing, 73% (CI 64 - 80%) and 76% (CI 67 – 82%), respectively. Both etofenprox-treated ranger uniform and civilian clothing continued to provide substantial protection after 20 washes, 65% (CI 53 - 74%) and 61% (CI 48 - 71%), respectively (Fig. 4, Table 3).

**Table 3.** Estimated rate ratios (RR) and their derived protective efficacies by intervention.

| Intervention | RR | 95% CI | p-value | Protective efficacy (%)<br>(%): 100 x (1-RR) | PE 95% CI (%) |
| --- | --- | --- | --- | --- | --- |
| BB+EtoC-0-Pi | 0.05 | [0.04 – 0.07] | <0.001 | 95% | [93 – 96] |
| BB | 0.06 | [0.05 – 0.09] | <0.001 | 94% | [91 – 95] |
| EtoC-0-Pi | 0.24 | [0.18 – 0.33] | <0.001 | 76% | [67 – 82] |
| EtoC-20-Pi | 0.39 | [0.29 – 0.52] | <0.001 | 61% | [48 – 71] |
| EtoR-0-Pi | 0.27 | [0.20 – 0.36] | <0.001 | 73% | [64 – 80] |
| EtoR-20-Pi | 0.35 | [0.26 – 0.47] | <0.001 | 65% | [53 – 74] |

**Fig. 3.**
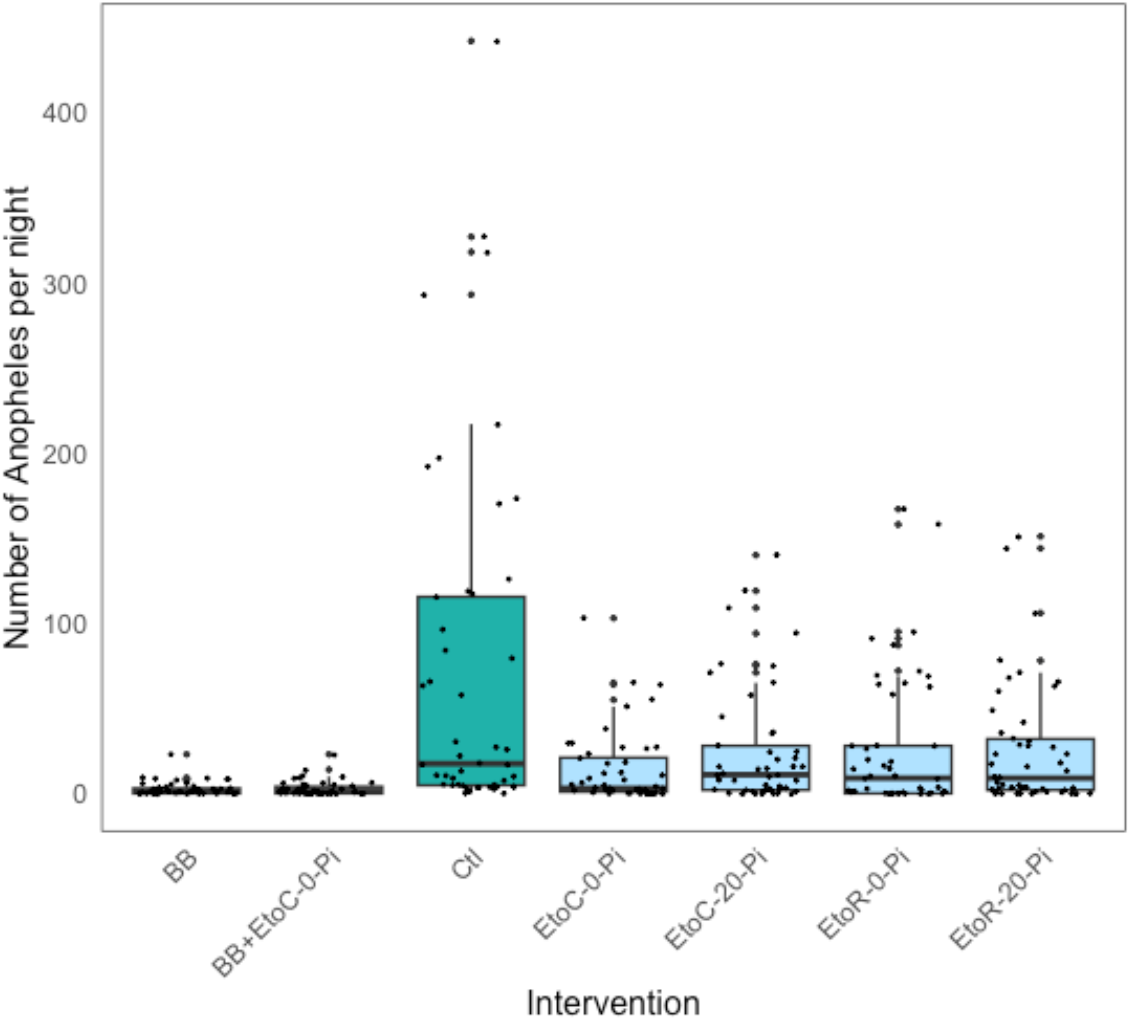
Number of *Anopheles* captured per HLC night by intervention.

**Fig. 4.**
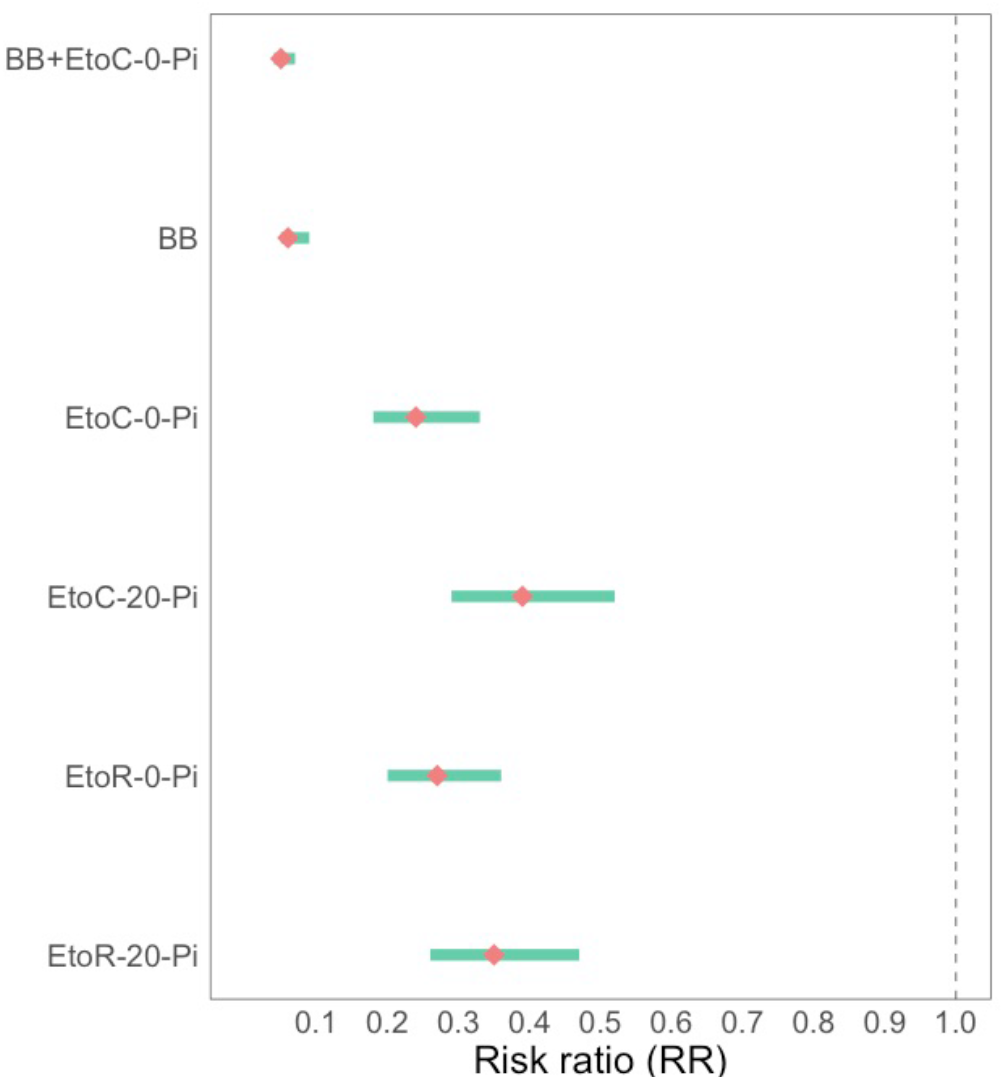
Estimated risk ratios (95% CI) for the effect of each intervention on risk of *Anopheles* landing (plotted on log scale).

Overall, there was variation in the number of *Anopheles* captured per night by both the structure locations (Suppl Fig. 1), and collector pair (Suppl Fig. 2). Environmental parameters (temperature, %RH, wind speed and rainfall) were examined against total *Anopheles* captures per night. Temperatures remained stable, between 22ºC and 24ºC, and wind speed did not exceed 1.2 m/s; there was no clear relationship between temperatures or wind speed with *Anopheles* captures. Humidity was highest (100%) during the period of heaviest rainfalls (October), which coincided with lower *Anopheles* catches.

### Relative reduction in vectoral capacity

This parameter was measured by combining field data and the semi-field data generated during the preceding SFS trials in Thailand were to predict the potential of the tools to provide community protection. The relative reduction in vectoral capacity for each intervention coverage is shown in Figure 5. Overall, all interventions were predicted to have a large impact on the vectoral capacity. The impact increases with coverage and adherence. In line with the reduction in landing observed in the field study, the greatest reduction of vectoral capacity was predicted for the combined EtoC and BB VPSR as well as the BB VPSR used alone. Even at 50% usage large reductions in vectoral capacity are predicted. As interventions age, they are less effective at reducing the vectoral capacity (Figure 5).

**Fig. 5.**
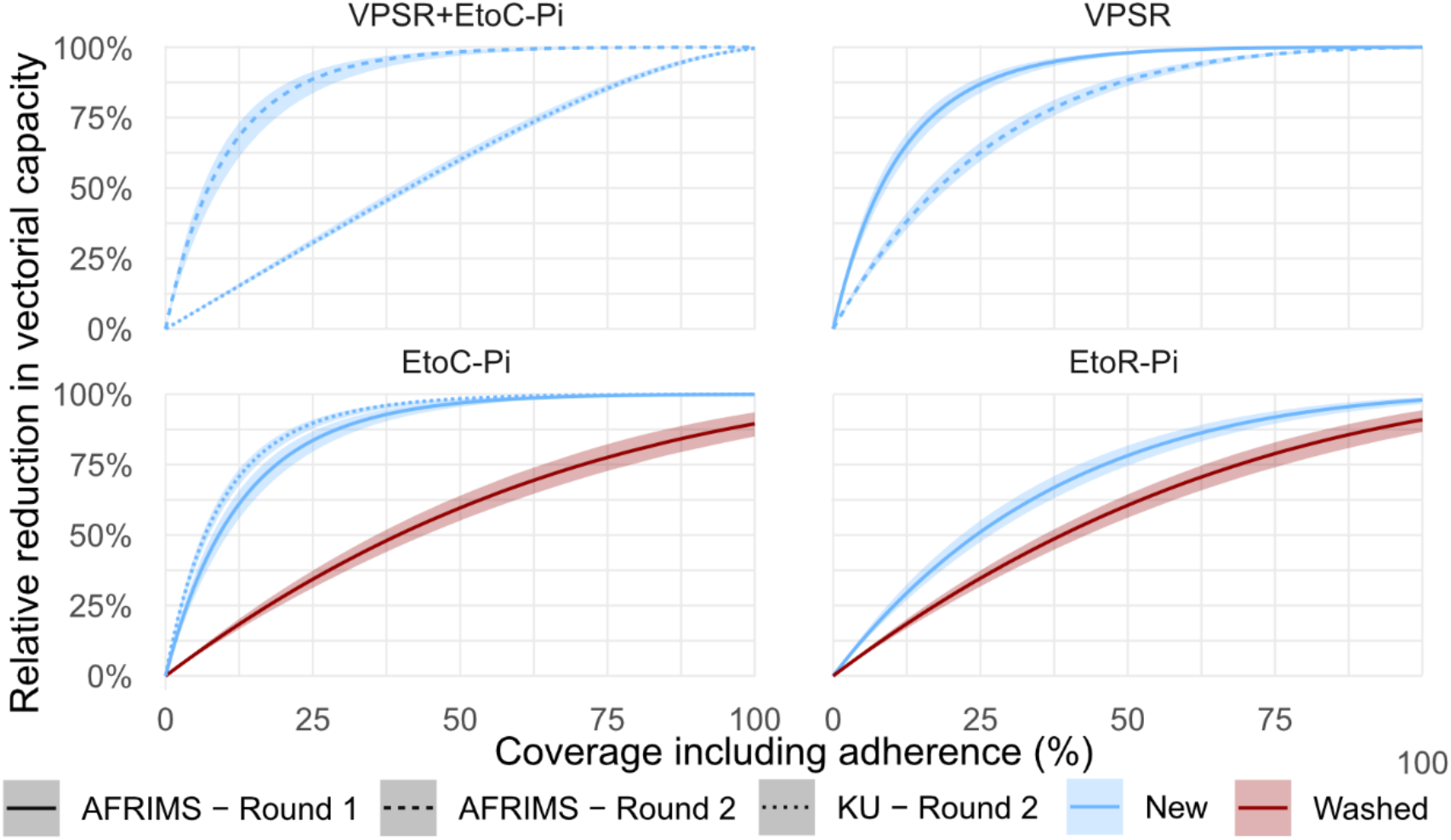
Predicted relative reduction in vectoral capacity for *Anopheles dirus* due to each intervention for a range of coverage levels for each intervention trailed in the field experiments. For interventions with semi-field estimates from multiple sites and years an individual line is plotted for each semi-field scenario. Black lines represent the median estimates, and colored bands represent the 95% CIs.

Although predictions have similar trends when utilizing data from different semi-field experiments, the magnitude of the predicted reduction in vectoral capacity is variable. There were differences in the extent of the modes of action observed, and therefore the parameter estimates, between experiments. These could be due to a number of factors including meteorological differences between locations and times and genetic differences between mosquito colonies.

## Discussion

In this field evaluation, the authors measured the impact of outdoor use of a passive, transfluthrin-based spatial repellent and etofenprox-treated clothing (paired with a 20% picaridin topical repellent) in the form of six bite prevention interventions on wild *Anopheles* landing. The study was designed to estimate landing rates (as a proxy for biting rates) when using the interventions while sitting in a fixed position in open structures similar to the open shelters used by forest-exposed populations in Cambodia.

A wide diversity of *Anopheles* species was observed in the study site in Mondulkiri (Table 2). Of the 15 *Anopheles* species collected, five are known vectors of human malaria: *An. dirus* Form A [58,59,12], *An. maculatus* [60,61], *An. minimus* [61,62], *An. sawadwongporni* [61], and *An. aconitus* [7,63]. Species *An. dirus, An. maculatus*, and *An. minimus* comprise Cambodia’s major malaria vectors [5,64,65], and all 15 collected species have been previously identified in Cambodia [7,61,66]. While HLC collections occurred from early evening (18h00) to early morning (06h00), it is possible that daytime biting would also be observed had collections taken place prior to 18h00. Two other entomological studies in Mondulkiri province, one being in a forested site [4], and the other being in both forested and village sites [7], included 24-hour *Anopheles* collections demonstrating daytime biting. Daytime biting suggests a further need for additional bite prevention interventions that are protective even when users are awake and outdoors [4,7].

This field study found that all of the study interventions, even when washed (for the treated clothing interventions), provided substantial protection against wild *Anopheles* landing outdoors in open temporary structures (Table 3, Fig. 6). The BB VPSR was highly effective against *Anopheles* landing, preventing 94% (CI 91 - 95%) of landings (Table 2). Combining this passive VPSR with etofenprox-treated clothing made no substantial difference on the level of protection provided against *Anopheles* landing. However, the combined intervention – delivered as a forest pack – is intended to protect real-world users both while inside their homes or temporary shelters and while mobile (outside their homes/shelters), offering “full-time” or 24-hour protection. In other words, the interventions are not meant to all be used at once but rather as needed based on individuals’ daily activities. For this study, the HLC collectors were restricted to the temporary structures.

Etofenprox-treated clothing interventions also provided high levels of protection against mosquito landing, though with slightly lower estimated protective efficacies than the BB VPSR intervention. However, while landing rates are often used as a proxy for measuring biting rates [55], evidence indicates that landing might not always necessarily lead to biting. When mosquitoes are exposed to pyrethroids (airborne or applied to clothing), they may still be able to detect the host and land, but they are inhibited from biting [67]. In a lab study, the landing and biting rates of *Aedes aegypti* were measured while using metofluthrin VPSRs, and observed 74 landings for only eight bites [68]. Therefore, it cannot be excluded that measuring landing rates may lead to an overestimation of ‘biting’ and an under estimation of protection. To more accurately estimate the protective efficacy of ITCs it might be best to allow mosquitoes to feed under controlled SFS conditions [69].

This study only evaluated etofenprox-treated clothing paired with a topical repellent, as the preceding SFS trials demonstrated that the combined ITC and topical repellent intervention was more effective against mosquito landing than the treated clothing alone (Vajda et al., unpublished 2020/2021 data, manuscript underway). In another field study evaluating personal repellent and treated clothing interventions in Lao PDR, permethrin-treated short clothing paired with a topical repellent were found to provide substantially more protection against mosquito landing than the permethrin-treated short clothing alone [55]. However, the permethrin-treated *long* clothing provided much higher protection against biting than the treated *short* clothing, but treated long clothing with a topical repellent was not tested [55]. Given the challenges with adherence around topical repellents use [17,70,71], it would be useful to compare treated long clothing alone, to treated long clothing with a topical repellent. This would help better understand how much more protection against biting is conferred by the addition of a topical repellent.

On the other hand, this study included a negative control with *short* trousers, but not one with *long* trousers (due to resource and time constraints). However, because the etofenprox-treated clothing interventions comprised long trousers that were *not* rolled up to the knees during HLCs, it is possible that the ratios of *Anopheles* landing in the treated clothing interventions compared to the *Anopheles* landing in the control slightly overestimate the landing protective efficacy since the negative control did not have the added physical barrier provided by ankle-length trousers.

Washing the etofenprox-treated clothing interventions 20 times only slightly increased the risk of *Anopheles* landing. This finding has implications for determining the frequency of retreatment of etofenprox-treated clothing. Efficacy testing of ITCs in controlled field and/or laboratory settings should be regarded as a proxy for real-world field conditions, in which ITCs would face harsher and more variable conditions (e.g., intense sweating during physical labor, textile degradation). Therefore, care should be taken when interpreting controlled testing results, and where possible, test results should be considered alongside results from efficacy testing in real-world field settings as is conducted for other vector control tools such as ITNs [72].

Permethrin is the most common insecticide used to treat clothing [73]. Given the shortcomings of permethrin regarding longevity, efficacy on lightweight fabrics, and potential carcinogenicity, etofenprox formulations for clothing treatment have been developed and registered by the U.S. Environmental Protection Agency [74]. Etofenprox is a synthetic pyrethroid-like ether insecticide. The structure of etofenprox renders it more stable and less toxic than other pyrethroid insecticides and functions by attacking the neuronal axon of the mosquito. As concerns over the spread of pyrethroid resistance grows, and several studies of the impact of pyrethroid resistance on the protective effect treated clothing yield conflicting results [75,34,76,35] there is growing need to explore alternative insecticides for clothing treatment.

The set replacement rate of the BB VPSR in the field must also be based on its residual efficacy. This study only tested new, freshly manufactured BB products towards providing a baseline understanding of the product’s efficacy against local, wild *Anopheles*, and did not investigate its residual efficacy. Therefore, the BB’s residual efficacy is currently unknown. Similarly to ITCs, transfluthrin-based VPSR efficacy over time is dependent on exposure to UV and other environmental parameters (rainfall, wind), as well as the susceptibility of local mosquitoes to pyrethroids [18,19,56,56,68,77]. For this field study, insecticide susceptibility testing to pyrethroids, including etofenprox and transfluthrin, was not feasible due to issues with keeping field-caught adult *Anopheles* alive and other logistical challenges.

This intervention evaluation provides evidence on the efficacy of these bite prevention interventions under controlled field conditions against wild *Anopheles* landing in Mondulkiri Province, Cambodia and highlights their potential for use in this elimination scenario if effective coverage of at-risk populations can be achieved. As with all vector control interventions, effectiveness is dependent on local vector bionomics [78], vector insecticide susceptibility profiles and resistance mechanisms [79–81], which uniquely affect intervention functionality [82–89]. Ecological, and human behavioral factors are also essential components that impact intervention effectiveness [47,90]. Therefore, malaria control and elimination programs must tailor their strategies and mix of interventions based on generated country-specific evidence and unique circumstances.

To date, the WHO has not established a position statement regarding the applications of VPSRs, ITCs, and topical repellents in public health vector control. However, while refraining from establishing formal recommendations on ITCs and topical repellents applications, the WHO does suggest these interventions for personal protection and considers their use for high-risk groups who do not benefit from other vector control interventions [39,91]. Recent work on a stochastic transmission model based on time-stratified vector landing data from controlled experiments of transfluthrin-treated eave ribbons (a type of VPSR) found that in addition to *Anopheles* vector landing reduction (personal protection), transfluthrin-treated ribbons also killed and reduced blood feeding, causing important reductions in vectoral capacity, indicating its potential to offer community protection [44]. The modeling evaluation conducted for this field study’s interventions’ on vectoral capacity also indicates reductions in vectoral capacity, even at decreased intervention coverage and adherence (Fig. 5). These works corroborate findings from recent SFS experiments [18–22,24] and field studies of VPSRs [25,92,93]. Given this growing body of evidence, repellents are increasingly recognized as having high potential for public health use, but further evidence of epidemiological impact is needed for WHO to establish a policy recommendation for these interventions [94].

## Conclusion

In Southeast Asia, *Anopheles* bites occurring outside the protection of the traditional, homestead-centric interventions (IRS, LLINs), constitute important gaps in protection that call for novel bite prevention interventions. This field study is highly encouraging as it demonstrates that this transfluthrin-based BB VPSR and etofenprox ITCs provide substantial protection against *Anopheles* landing when used outside. Further, the study’s modeling analysis predicted that these interventions would reduce vectoral capacity, even at 50% coverage. Additional studies in other geographic settings are needed to estimate intervention impact on landing of mosquitoes with different bionomics profiles, and trials in which the products are distributed and used normally are also required. While further evidence of epidemiological impact is necessary for WHO to establish these tools as effective public health interventions against malaria, this study provides promising results toward addressing gaps in protection of at-risk groups against *Anopheles* biting.

## Acknowledgements

This study was funded by the Australia Department of Foreign Affairs and Trade (DFAT) through the Innovative Vector Control Consortium (IVCC), Liverpool, United Kingdom. We would also like to acknowledge and thank all the project teams, entomological collectors and technicians, local authorities, and fieldworkers who provided unwavering support throughout the study, even during the COVID-19 pandemic.

## Author contributions

EAV substantially contributed to study conceptualization and design, led writing of protocol, SOPs, and data collection tools, led data curation, analysis, and synthesis, and led manuscript writing and review process. AR substantially contributed to protocol writing, data analysis, and manuscript review. DD led all fieldwork activities and contributed to manuscript review. EF led the modeling component and write-up in the manuscript, and also substantially contributed to manuscript review. NC supported the modeling component of the paper and contributed to manuscript review. JH substantially contributed to study design, protocol, and SOP writing, fieldwork activities, and manuscript review. SJM substantially contributed to study conceptualization, protocol writing, data synthesis, and manuscript review. JR contributed to manuscript review. MM contributed contributed to manuscript review. SS contributed to study conceptualization and facilitated study rollout in Cambodia. PK provided support to fieldwork planning and activities. DJM substantially contributed to study conceptualization, design, and protocol writing, and manuscript review. AT substantially contributed to study conceptualization, design, and protocol writing, and manuscript review. NFL substantially contributed to study conceptualization, protocol writing, fieldwork, and manuscript review.

## Data availability

Correspondence and requests for data should be directed to EAV.

## Additional information

### Supplementary information

*See below*

## Supplementary materials

**Table 1.** Definitions and values of parameters used to calculate the vectoral capacity. Baseline values, considering all humans as unprotected, for *Plasmodium falciparum* malaria and *Anopheles dirus* are given.

| Definition | Value | Ref. |
| --- | --- | --- |
| Number of humans | 10,000 | Assumption |
| Number of non-human hosts | 10,000 | Assumption |
| Maximum time a mosquito unsuccessfully searches for a blood meal per day | 0.33 days | Briët et al. 2019 [95] |
| Probability that a mosquito bites after encountering a host | 0.95 | Briët et al. 2019 |
| Probability that a mosquito finds a resting place | 0.95 | Briët et al. 2019 |
| Probability that a mosquito survives the resting phase | 0.99 | Briët et al. 2019 |
| Probability that a mosquito lays eggs and returns to host-seeking | 0.88 | Briët et al. 2019 |
| Time between feeding and laying eggs | 3 days | Wang et.al. (preprint) 2023 [96] |
| Duration of the extrinsic incubation period (time required for sporozoites to develop in the mosquito) | 10 days | Briët et al. 2019 |
| The sac rate of mosquitoes (estimated proportion of host-seeking mosquitoes which laid eggs the same day) | 0.33 | Wang et.al. (preprint) 2023 |
| The parity rate of mosquitoes (proportion of host-seeking mosquitoes that have previously laid eggs) | 0.64 | Wang et.al. (preprint) 2023 |
| Human blood index (proportion of blood-fed mosquitoes which fed on a human) | 0.87 | Wang et.al. (preprint) 2023 |
| Human availability rate (the rate at which mosquitoes encounter humans) | $8.51 \times 10^{-5} \text{ days}^{-1}$ | Calculated (as described in Briët et al. 2019) |
| Non-human availability rate (the rate at which mosquitoes encounter animals) | $1.27 \times 10^{-4} \text{ days}^{-1}$ | Calculated (as described in Briët et al. 2019) |
| Per-capita mosquito death rate while searching for a blood meal | $0.22 \text{ days}^{-1}$ | Calculated (as described in Briët et al. 2019) |

**suppl Fig. 1.**
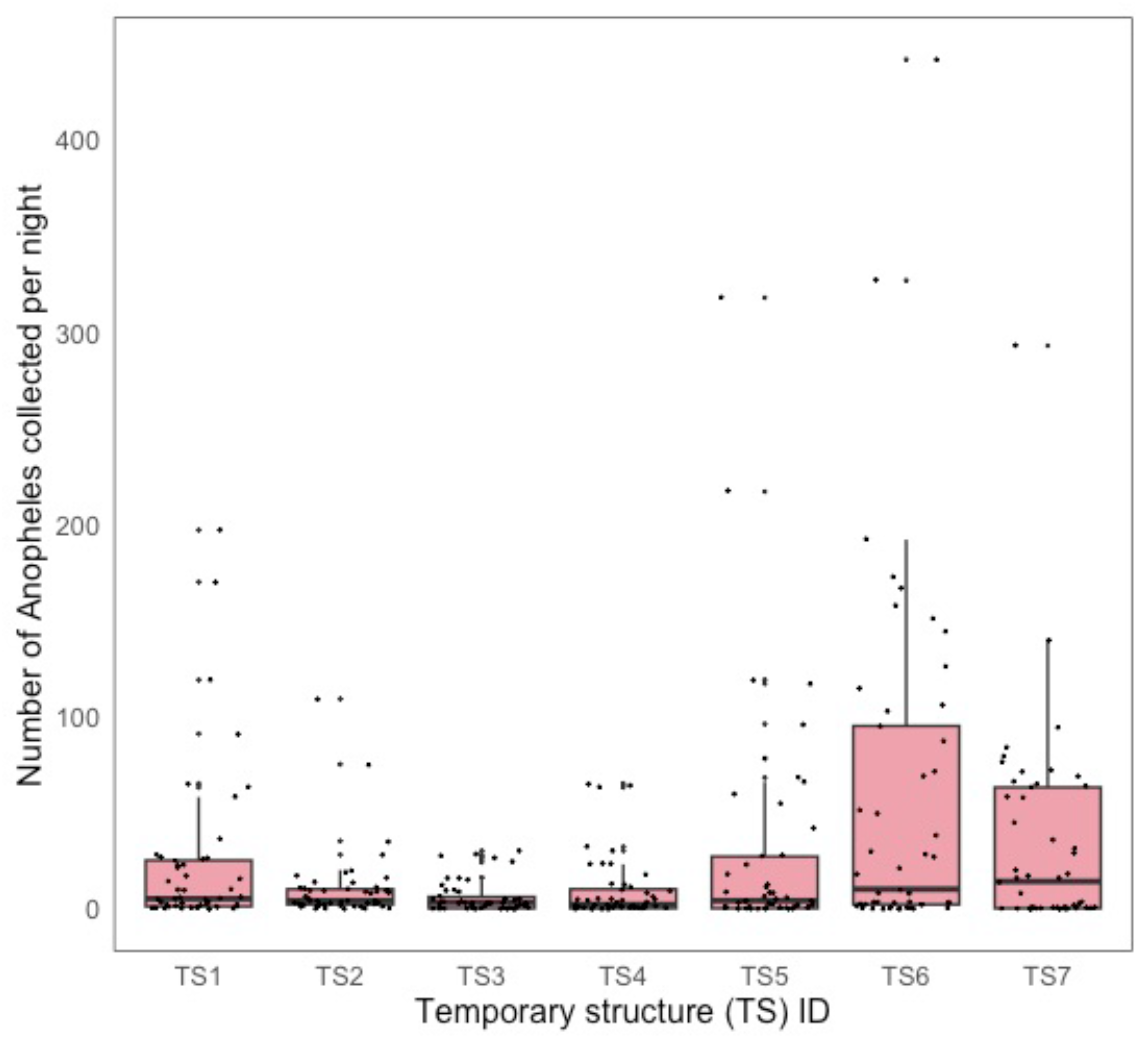
*Anopheles* captured per HLC night, per temporary structure location.

**suppl Fig. 2.**
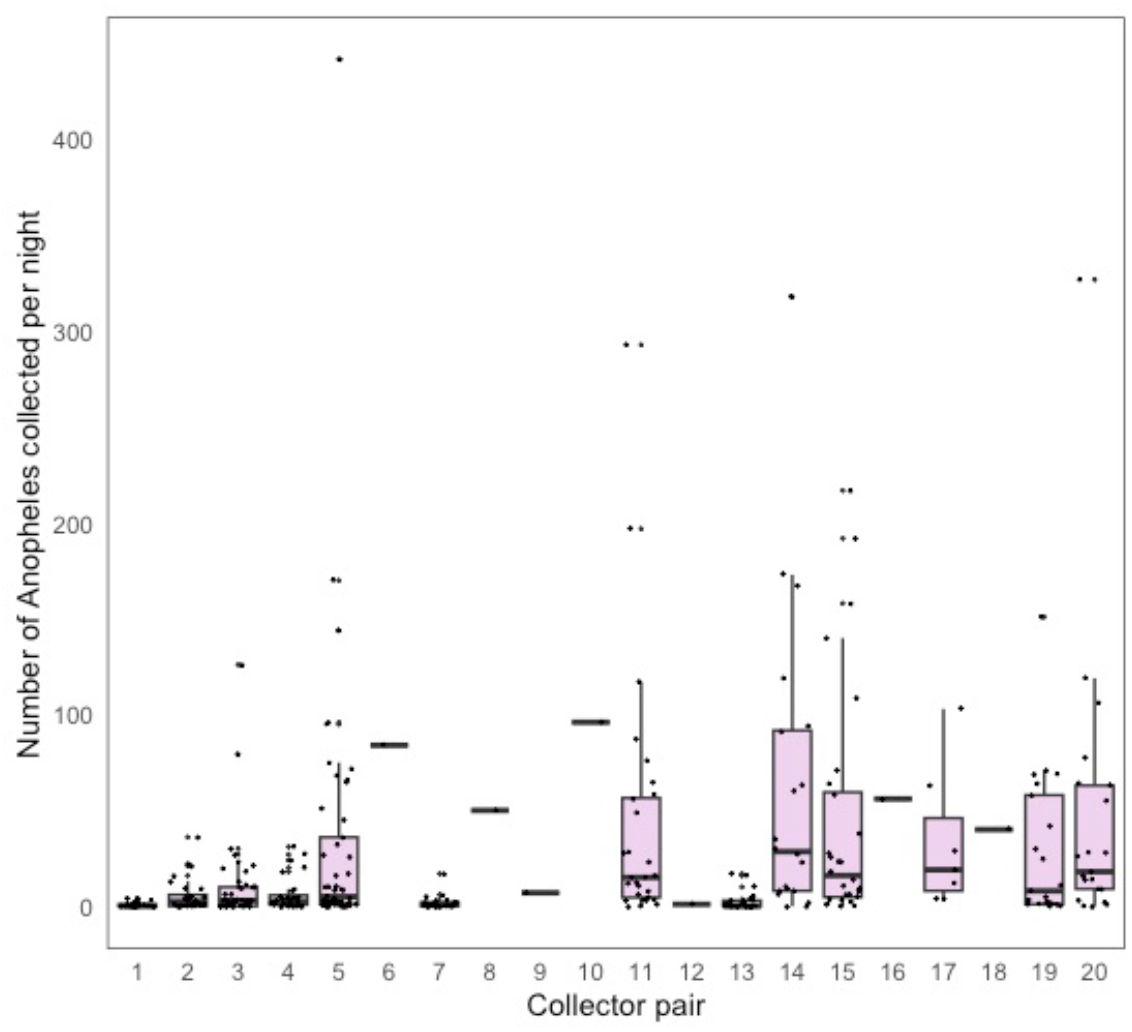
*Anopheles* captured per HLC night, per collector pair.

## References

1. World Health Organization. World malaria report 2023. Geneva: World Health Organization; 2023.

2. Sovannaroth S, Ngor P, Khy V, Dunn JC, Burbach MK, Peng S, et al. Accelerating malaria elimination in Cambodia: an intensified approach for targeting at-risk populations. Malar J. 2022;21:209.

3. Durnez L, Mao S, Denis L, Roelants P, Sochantha T, Coosemans M. Outdoor malaria transmission in forested villages of Cambodia. Malar J. 2013;12:329.

4. Kunkel A, Nguon C, Iv S, Chhim S, Peov D, Kong P, et al. Choosing interventions to eliminate forest malaria: preliminary results of two operational research studies inside Cambodian forests. Malar J. 2021;20:51.

5. Cui L, Yan G, Sattabongkot J, Cao Y, Chen B, Chen X, et al. Malaria in the Greater Mekong Subregion: Heterogeneity and complexity. Acta Tropica. 2012;121:227–39.

6. Canavati SE, Kelly GC, Quintero CE, Vo TH, Tran LK, Ngo TD, et al. Targeting high risk forest goers for malaria elimination: a novel approach for investigating forest malaria to inform program intervention in Vietnam. BMC Infect Dis. 2020;20:757.

7. Vantaux A, Riehle MM, Piv E, Farley EJ, Chy S, Kim S, et al. Anopheles ecology, genetics and malaria transmission in northern Cambodia. Sci Rep. 2021;11:6458.

8. U.S. President’s Malaria Initiative. U.S. President’s Malaria Initiative Cambodia Malaria Operational Plan FY 2022. 2021.

9. National Center for Parasitology, Entomology, and Malaria Control. National Treatment Guidelines for Malaria in Cambodia. 2022.

10. World Health Organization. Countries of the Greater Mekong ready for the “last mile” of malaria elimination, BUlletin #9. 2020.

11. Van--Roey K, Sokny M, Denis L, Van--den--Broeck N, Heng S, Siv S, et al. Field Evaluation of Picaridin Repellents Reveals Differences in Repellent Sensitivity between Southeast Asian Vectors of Malaria and Arboviruses. Vosshall L, editor. PLoS Negl Trop Dis. 2014;8:e3326.

12. St. Laurent B, Oy K, Miller B, Gasteiger EB, Lee E, Sovannaroth S, et al. Cow-baited tents are highly effective in sampling diverse Anopheles malaria vectors in Cambodia. Malar J. 2016;15:440.

13. Sanann N, Peto TJ, Tripura R, Callery JJ, Nguon C, Bui TM, et al. Forest work and its implications for malaria elimination: a qualitative study. Malar J. 2019;18:376.

14. Bibbs CS, Kaufman PE. Volatile Pyrethroids as a Potential Mosquito Abatement Tool: A Review of Pyrethroid-Containing Spatial Repellents. Journal of Integrated Pest Management [Internet]. 2017 [cited 2022 Feb 13];8. Available from: 10.1093/jipm/pmx016

15. Kongmee M, Boonyuan W, Achee NL, Prabaripai A, Lerdthusnee K, Chareonviriyaphap T. Irritant and Repellent Responses of Anopheles harrisoni and Anopheles minimus upon Exposure to Bifenthrin or Deltamethrin Using an Excito-Repellency System and a Live Host. Journal of the American Mosquito Control Association. 2012;28:20–9.

16. Banks SD, Murray N, Wilder-Smith A, Logan JG. Insecticide-treated clothes for the control of vector-borne diseases: a review on effectiveness and safety. Med Vet Entomol. 2014;28:14–25.

17. Maia MF, Kliner M, Richardson M, Lengeler C, Moore SJ. Mosquito repellents for malaria prevention. Cochrane Infectious Diseases Group, editor. Cochrane Database of Systematic Reviews [Internet]. 2018 [cited 2022 Dec 21];2018. Available from: 10.1002/14651858.CD011595.pub2

18. Ogoma SB, Mmando AS, Swai JK, Horstmann S, Malone D, Killeen GF. A low technology emanator treated with the volatile pyrethroid transfluthrin confers long term protection against outdoor biting vectors of lymphatic filariasis, arboviruses and malaria. PLoS Negl Trop Dis. 2017;11:e0005455.

19. Masalu JP, Finda M, Okumu FO, Minja EG, Mmbando AS, Sikulu-Lord MT, et al. Efficacy and user acceptability of transfluthrin-treated sisal and hessian decorations for protecting against mosquito bites in outdoor bars. Parasit Vectors. 2017;10:197.

20. Stevenson JC, Simubali L, Mudenda T, Cardol E, Bernier UR, Vazquez AA, et al. Controlled release spatial repellent devices (CRDs) as novel tools against malaria transmission: a semi-field study in Macha, Zambia. Malar J. 2018;17:437.

21. Tambwe MM, Moore SJ, Chilumba H, Swai JK, Moore JD, Stica C, et al. Semi-field evaluation of freestanding transfluthrin passive emanators and the BG sentinel trap as a “push-pull control strategy” against Aedes aegypti mosquitoes. Parasites Vectors. 2020;13:392.

22. Sangoro OP, Gavana T, Finda M, Mponzi W, Hape E, Limwagu A, et al. Evaluation of personal protection afforded by repellent-treated sandals against mosquito bites in south-eastern Tanzania. Malar J. 2020;19:148.

23. Tambwe MM, Moore S, Hofer L, Kibondo UA, Saddler A. Transfluthrin eave-positioned targeted insecticide (EPTI) reduces human landing rate (HLR) of pyrethroid resistant and susceptible malaria vectors in a semi-field simulated peridomestic space. Malar J. 2021;20:357.

24. Burton TA, Kabinga LH, Simubali L, Hayre Q, Moore SJ, Stevenson JC, et al. Semi-field evaluation of a volatile transfluthrin-based intervention reveals efficacy as a spatial repellent and evidence of other modes of action. De--Souza DK, editor. PLoS ONE. 2023;18:e0285501.

25. Sukkanon C, Tisgratog R, Muenworn V, Bangs MJ, Hii J, Chareonviriyaphap T. Field Evaluation of a Spatial Repellent Emanation Vest for Personal Protection Against Outdoor Biting Mosquitoes. Lysyk T, editor. Journal of Medical Entomology. 2021;58:756–66.

26. Charlwood JD, Nenhep S, Protopopoff N, Sovannaroth S, Morgan JC, Hemingway J. Effects of the spatial repellent metofluthrin on landing rates of outdoor biting anophelines in Cambodia, Southeast Asia: Metofluthrin in Cambodia. Med Vet Entomol. 2016;30:229–34.

27. Martin JA, Hendershot AL, Saá Portilla IA, English DJ, Woodruff M, Vera-Arias CA, et al. Anopheline and human drivers of malaria risk in northern coastal, Ecuador: a pilot study. Malar J. 2020;19:354.

28. Ponlawat A, Kankaew P, Chanaimongkol S, Pongsiri A, Richardson JH, Evans BP. Semi-Field Evaluation of Metofluthrin-Impregnated Nets on Host-Seeking Aedes aegypti and Anopheles dirus. Journal of the American Mosquito Control Association. 2016;32:130–8.

29. Yan C, Hii J, Ngoen-Klan R, Saeung M, Chareonviriyaphap T. Semi-field evaluation of human landing catches versus human double net trap for estimating human biting rate of Anopheles minimus and Anopheles harrisoni in Thailand. PeerJ. 2022;10:e13865.

30. Pennetier C, Chabi J, Martin T, Chandre F, Rogier C, Hougard J-M, et al. New protective battle-dress impregnated against mosquito vector bites. Parasites Vectors. 2010;3:81.

31. Most B, de Santi VP, Pagès F, Mura M, Uedelhoven WM, Faulde MK. Long-lasting permethrin-impregnated clothing: protective efficacy against malaria in hyperendemic foci, and laundering, wearing, and weathering effects on residual bioactivity after worst-case use in the rain forests of French Guiana. Parasitol Res. 2017;116:677–84.

32. Williams YA, Tusting LS, Hocini S, Graves PM, Killeen GF, Kleinschmidt I, et al. Expanding the vector control toolbox for malaria elimination: a systematic review of the evidence. Adv Parasitol. 2018;99:345–79.

33. Vajda ÉA, Saeung M, Ross A, McIver DJ, Tatarsky A, Moore SJ, et al. A semi-field evaluation in Thailand of the use of human landing catches (HLC) versus human-baited double net trap (HDN) for assessing the impact of a volatile pyrethroid spatial repellent and pyrethroid-treated clothing on Anopheles minimus landing. Malar J. 2023;22:202.

34. Gordon, Scott W. Personal protection tools from the deployed warfighter research program (DWFP) [Internet]. 2013. Available from: https://endmalaria.org/sites/default/files/4_Scott%20Gordon_DWFP_Outdoor%20WS.pdf

35. Combating the Aedes aegypti [Internet]. perimiterinsectguard.com/eto/. Available from: https://www.perimeterinsectguard.com/eto/

36. Eamsila C, Frances SP. EVALUATION OF PERMETHRIN.TREATED MILITARY UNIFORMS FOR PERSONAL PROTECTION AGAINST MALARIA IN NORTHEASTERN THAILAND’.

37. Rowland M, Durrani N, Hewitt S, Mohammed N, Bouma M, Carneiro I, et al. Permethrin-treated chaddars and top-sheets: appropriate technology for protection against malaria in Afghanistan and other complex emergencies. Transactions of the Royal Society of Tropical Medicine and Hygiene. 1999;93:465–72.

38. Kittayapong P, Olanratmanee P, Maskhao P, Byass P, Logan J, Tozan Y, et al. Mitigating Diseases Transmitted by Aedes Mosquitoes: A Cluster-Randomised Trial of Permethrin-Impregnated School Uniforms. Apperson C, editor. PLoS Negl Trop Dis. 2017;11:e0005197.

39. World Health Organization. WHO guidelines for malaria, 16 October 2023. Geneva: World Health Organization; 2023.

40. Vontas J, Moore S, Kleinschmidt I, Ranson H, Lindsay S, Lengeler C, et al. Framework for rapid assessment and adoption of new vector control tools. Trends in Parasitology. 2014;30:191–204.

41. Garros C, Van Bortel W, Trung HD, Coosemans M, Manguin S. Review of the Minimus Complex of Anopheles, main malaria vector in Southeast Asia: from taxonomic issues to vector control strategies. Trop Med Int Health. 2006;11:102–14.

42. Fairbanks EL, Saeung M, Pongsiri A, Vajda E, Wang Y, McIver DJ, et al. Inference for entomological semi-field experiments: Fitting a mathematical model assessing personal and community protection of vector-control interventions [Internet]. Systems Biology; 2023 Jul. Available from: 10.1101/2023.06.30.547223

43. Garrett-Jones C, Shidrawi GR. Malaria Vectorial Capacity of a Population of Anopheles ganmbiae.

44. Denz A, Njoroge MM, Tambwe MM, Champagne C, Okumu F, van--Loon JJA, et al. Predicting the impact of outdoor vector control interventions on malaria transmission intensity from semi-field studies. Parasites Vectors. 2021;14:64.

45. Miyamoto J. Degradation, Metabolism and Toxicity of Synthetic Pyrethroids. Environmental Health Perspectives. 1976;

46. Ogoma SB, Ngonyani H, Simfukwe ET, Mseka A, Moore J, Maia MF, et al. The Mode of Action of Spatial Repellents and Their Impact on Vectorial Capacity of Anopheles gambiae sensu stricto. Hansen IA, editor. PLoS ONE. 2014;9:e110433.

47. Chen I, Doum D, Mannion K, Hustedt J, Sovannaroth S, McIver D, et al. Applying the COM-B behaviour change model to a pilot study delivering volatile pyrethroid spatial repellents and insecticide-treated clothing to forest-exposed populations in Mondulkiri Province, Cambodia. Malar J. 2023;22:251.

48. Rattanarithikul R, Harrison BA, Harbach RE, Panthusiri P, Coleman RE. ILLUSTRATED KEYS TO THE MOSQUITOES OF THAILAND IV. Anopheles.

49. www.boldsystems.org. Barcode of Life Data Systems Handbook: A web-based bioinformatics platform supporting the DNA barcoding of animal, plant, and fungal species. 2023.

50. R Core Team. R: A language and environment for statistical computing. Vienna, Austria; 2022.

51. Hadley Wickham, Davis Vaughan and Maximilian Girlich. tidyr: Tidy Messy Data. R package version 1.3.0. [Internet]. 2023. Available from: https://CRAN.R-project.org/package=tidyr

52. Hadley Wickham, Romain François, Lionel Henry and Kirill Müller. dplyr: A Grammar of Data Manipulation. R package version 1.0.8. [Internet]. 2022. Available from: https://CRAN.R-project.org/package=dplyr

53. Bates D, Mächler M, Bolker B, Walker S. Fitting Linear Mixed-Effects Models Using lme4. J Stat Soft [Internet]. 2015 [cited 2023 Jul 27];67. Available from: http://www.jstatsoft.org/v67/i01/

54. Wickham H. ggplot2: Elegant Graphics for Data Analysis. Springer-Verlag New York; 2016.

55. Tangena J-AA, Thammavong P, Chonephetsarath S, Logan JG, Brey PT, Lindsay SW. Field evaluation of personal protection methods against outdoor-biting mosquitoes in Lao PDR. Parasites Vectors. 2018;11:661.

56. Tambwe MM, Kibondo UA, Odufuwa OG, Moore J, Mpelepele A, Mashauri R, et al. Human landing catches provide a useful measure of protective efficacy for the evaluation of volatile pyrethroid spatial repellents. Parasites Vectors. 2023;16:90.

57. Fassler J, Cooper P. BLAST Glossary. 2011 Jul 14. In: BLAST® Help [Internet]. Bethesda (MD): National Center for Biotechnology Information (US); 2008.

58. Peyton EL, Harrison BA. Anopheles (Celia) dirus, a New Species of the Leucosphyrus Group from Thailand (Diptera: Culicidae). 1979;11.

59. Manguin S, Garros C, Dusfour I, Harbach RE, Coosemans M. Bionomics, taxonomy, and distribution of the major malaria vector taxa of Anopheles subgenus Cellia in Southeast Asia: An updated review. Infection, Genetics and Evolution. 2008;8:489–503.

60. Rahman WA, Hassan AA, Adanan CR, Rashid MohdR. The prevalence of Plasmodium falciparum and P. vivax in relation to Anopheles maculatus densities in a Malaysian village. Acta Tropica. 1993;55:231–5.

61. Sinka ME, Bangs MJ, Manguin S, Chareonviriyaphap T, Patil AP, Temperley WH, et al. The dominant Anopheles vectors of human malaria in the Asia-Pacific region: occurrence data, distribution maps and bionomic précis. Parasites Vectors. 2011;4:89.

62. Tananchai C, Manguin S, Bangs MJ, Chareonviriyaphap T. Malaria Vectors and Species Complexes in Thailand: Implications for Vector Control. Trends in Parasitology. 2019;35:544–58.

63. Chen B, Harbach RE, Walton C, He Z, Zhong D, Yan G, et al. Population genetics of the malaria vector Anopheles aconitus in China and Southeast Asia. Infection, Genetics and Evolution. 2012;12:1958–67.

64. Vantaux A, Samreth R, Piv E, Khim N, Kim S, Berne L, et al. Contribution to Malaria Transmission of Symptomatic and Asymptomatic Parasite Carriers in Cambodia. The Journal of Infectious Diseases. 2018;217:1561–8.

65. Zhang C, Yang R, Wu L, Luo C, Yang Y, Deng Y, et al. Survey of malaria vectors on the Cambodia, Thailand and China-Laos Borders. Malar J. 2022;21:399.

66. St. Laurent B, Oy K, Miller B, Gasteiger EB, Lee E, Sovannaroth S, et al. Cow-baited tents are highly effective in sampling diverse Anopheles malaria vectors in Cambodia. Malar J. 2016;15:440.

67. Lucas JR, Shono Y, Iwasaki T, Ishiwatari T, Spero N, Benzon G. U.S. LABORATORY AND FIELD TRIALS OF METOFLUTHRIN (SUMIONE®) EMANATORS FOR REDUCING MOSQUITO BITING OUTDOORS ^1^. Journal of the American Mosquito Control Association. 2007;23:47–54.

68. Mmbando AS, Ngowo HS, Kilalangongono M, Abbas S, Matowo NS, Moore SJ, et al. Small-scale field evaluation of push-pull system against early-and outdoor-biting malaria mosquitoes in an area of high pyrethroid resistance in Tanzania. Wellcome Open Res. 2017;2:112.

69. Swai JK, Kibondo UA, Ntabaliba WS, Ngoyani HA, Makungwa NO, Mseka AP, et al. CDC light traps underestimate the protective efficacy of an indoor spatial repellent against bites from wild Anopheles arabiensis mosquitoes in Tanzania. Malar J. 2023;22:141.

70. Wilson AL, Chen-Hussey V, Logan JG, Lindsay SW. Are topical insect repellents effective against malaria in endemic populations? A systematic review and meta-analysis. Malar J. 2014;13:446.

71. Sluydts V, Durnez L, Heng S, Gryseels C, Canier L, Kim S, et al. Efficacy of topical mosquito repellent (picaridin) plus long-lasting insecticidal nets versus long-lasting insecticidal nets alone for control of malaria: a cluster randomised controlled trial. The Lancet Infectious Diseases. 2016;16:1169–77.

72. World Health Organization. Guidelines for monitoring the durability of long-lasting insecticidal mosquito nets under operational conditions. 2011;44.

73. Goodyer LI, Croft AM, Frances SP, Hill N, Moore SJ, Onyango SP, et al. Expert Review of the Evidence Base for Arthropod Bite Avoidance. J Travel Med. 2010;17:182–92.

74. U.S Environmental Protection Agency. Notice of pesticide registration [Internet]. 2016. Available from: https://www3.epa.gov/pesticides/chem_search/ppls/082392-00003-20160822.pdf

75. The Assistant Secretary of Defense, (Health Affairs). Updated Policy for Prevention of Arthropod-Borne Diseases Among Department of Defense Personnel Deployed to Endemic Areas [Internet]. Health Affairs Policy: 07-007; 2007. Available from: https://apps.dtic.mil/sti/pdfs/ADA512584.pdf

76. Estep AS, Sanscrainte ND, Cuba I, Allen GM, Becnel JJ, Linthicum KJ. Failure of Permethrin-Treated Military Uniforms to Protect Against a Laboratory-Maintained Knockdown-Resistant Strain of Aedes aegypti. Journal of the American Mosquito Control Association. 2020;36:127–30.

77. Andrés M, Lorenz LM, Mbeleya E, Moore SJ. Modified mosquito landing boxes dispensing transfluthrin provide effective protection against Anopheles arabiensis mosquitoes under simulated outdoor conditions in a semi-field system. Malar J. 2015;14:255.

78. Trung HD, Bortel WV, Sochantha T, Keokenchanh K, Briet OJT, Coosemans M. Behavioural heterogeneity of Anopheles species in ecologically different localities in Southeast Asia: a challenge for vector control. Trop Med Int Health. 2005;10:251–62.

79. Machani MG, Ochomo E, Amimo F, Mukabana WR, Githeko AK, Yan G, et al. Behavioral responses of pyrethroid resistant and susceptible Anopheles gambiae mosquitoes to insecticide treated bed net. Oliveira PL, editor. PLoS ONE. 2022;17:e0266420.

80. Hemingway J, Ranson H. Insecticide Resistance in Insect Vectors of Human Disease. Annu Rev Entomol. 2000;45:371–91.

81. Santolamazza F, Calzetta M, Etang J, Barrese E, Dia I, Caccone A, et al. Distribution of knock-down resistance mutations in Anopheles gambiae molecular forms in west and west-central Africa. Malar J. 2008;7:74.

82. Githeko AK, Service MW, Mbogo CM, Atieli FA, Juma FO. Sampling Anopheles arabiensis, A. gambiae sensu lato and A. funestus (Diptera: Culicidae) with CDC light-traps near a rice irrigation area and a sugarcane belt in western Kenya. Bull Entomol Res. 1994;84:319–24.

83. Chandre F, Darriet F, Duchon S, Finot L, Manguin S, Carnevale P, et al. Modifications of pyrethroid effects associated with kdr mutation in Anopheles gambiae. Med Vet Entomol. 2000;14:81–8.

84. Wagman JM, Achee NL, Grieco JP. Insensitivity to the Spatial Repellent Action of Transfluthrin in Aedes aegypti: A Heritable Trait Associated with Decreased Insecticide Susceptibility. Lenhart A, editor. PLoS Negl Trop Dis. 2015;9:e0003726.

85. Agramonte NM, Bloomquist JR, Bernier UR. Pyrethroid resistance alters the blood-feeding behavior in Puerto Rican Aedes aegypti mosquitoes exposed to treated fabric. Apperson C, editor. PLoS Negl Trop Dis. 2017;11:e0005954.

86. Bowman NM, Akialis K, Cave G, Barrera R, Apperson CS, Meshnick SR. Pyrethroid insecticides maintain repellent effect on knock-down resistant populations of Aedes aegypti mosquitoes. Vontas J, editor. PLoS ONE. 2018;13:e0196410.

87. Afify A, Betz JF, Riabinina O, Lahondère C, Potter CJ. Commonly Used Insect Repellents Hide Human Odors from Anopheles Mosquitoes. Current Biology. 2019;29:3669–3680.e5.

88. Sherrard-Smith E, Skarp JE, Beale AD, Fornadel C, Norris LC, Moore SJ, et al. Mosquito feeding behavior and how it influences residual malaria transmission across Africa. Proc Natl Acad Sci USA. 2019;116:15086–95.

89. Afify A, Potter CJ. Insect repellents mediate species-specific olfactory behaviours in mosquitoes. Malar J. 2020;19:127.

90. Liverani M, Charlwood JD, Lawford H, Yeung S. Field assessment of a novel spatial repellent for malaria control: a feasibility and acceptability study in Mondulkiri, Cambodia. Malar J. 2017;16:412.

91. World Health Organization. Data requirements and methods to support the evaluation of new vector control products. 2017;7.

92. Dia I, Diallo D, Duchemin J-B, Konate L, Costantini C, Diallo M. Comparisons of Human-Landing Catches and Odor-Baited Entry Traps for Sampling Malaria Vectors in Senegal. JOURNAL OF MEDICAL ENTOMOLOGY. 2005;42.

93. Charlwood JD, Rowland M, Protopopoff N, Le Clair C. The Furvela tent-trap Mk 1.1 for the collection of outdoor biting mosquitoes. PeerJ. 2017;5:e3848.

94. Achee NL, Perkins TA, Moore SM, Liu F, Sagara I, Van--Hulle S, et al. Spatial repellents: The current roadmap to global recommendation of spatial repellents for public health use. Current Research in Parasitology & Vector-Borne Diseases. 2023;3:100107.

95. Briët OJT, Impoinvil DE, Chitnis N, Pothin E, Lemoine JF, Frederic J, et al. Models of effectiveness of interventions against malaria transmitted by Anopheles albimanus. Malar J. 2019;18:263.

96. Wang Y, Chitnis N, Fairbanks E. Optimizing malaria vector control: A systematic review and mathematical modelling study to identify desirable characteristics of novel tools in different settings [Internet]. In Review; 2023 Sep. Available from: https://www.researchsquare.com/article/rs-3332552/v1

